# Intrinsic electrophysiological activity maps a latent dimension of poor sleep quality and reduced cognitive performance: a magnetoencephalography study using Cam-CAN data

**DOI:** 10.1101/2025.11.01.686036

**Authors:** Samuel Hardy, Gill Roberts, George O’Neill, Venkat Bhat, Yali Pan, Amy C Reichelt, Robert A Seymour, Benjamin T Dunkley

## Abstract

Sleep quality and cognition vary as functions of lifestyle, genetics, and health. However, poor sleep quality is prevalent, reportedly affecting approximately 38% of the adult population. Poor sleep quality is a significant risk factor for mood disorders, a predictive factor in cognitive decline in later life, and sleep disturbance is a putative precursor to severe cognitive impairment and is predictive of dementia onset. The direct relationship between these factors and intrinsic neural function is poorly understood. Integrating neurophysiology, cognition, and sleep quality to reveal latent factors would help understand the neurophysiology of sleep disturbances and their relation to cognitive performance. Here, we used data from the Cam-CAN dataset with a partial least squares (PLS) approach, producing a multivariate cross-decomposition model to map resting-state magnetoencephalography (MEG) data and cognitive/sleep scores of healthy controls (n = 490, age 18-86). Normative modelling was applied to MEG data, correcting for effects of age, sex, and handedness. We identify a significant relationship between poorer self-reported sleep quality and lower cognitive performance across multiple domains, characterised by excess low-frequency neural activity and reduced high-frequency activity, particularly in the alpha band. Moreover, globally increased low frequency and decreased alpha-beta functional connectivity – the dominant frequency channel in neural connectivity at rest across the brain – contribute significantly to this relationship. Our multivariate mapping approach enabled us to parse the electrophysiological signatures directly related to sleep quality and cognitive performance, independent of the confounding effects of age, and show that ‘neural slowing’ and dysconnectivity are intrinsically linked to self-reported sleep quality and cognitive performance.

## Introduction

Sleep and cognition are two critical components of daily life that have a profound impact on overall health and well-being. Sleep is not merely a period of rest but an essential biological function that plays a pivotal role in optimising cognitive processes such as memory consolidation (Klinzing et al., 2019), attention (Lim & Dinges, 2008), learning (Curcio et al., 2006), and executive functioning (Anderson et al., 2009). Yet, despite the importance of sleep, contemporary lifestyles have led to widespread sleep deprivation (SD) and poor sleep quality, exacerbated by factors such as work demands, digital screen exposure, and societal pressures (Shochat, 2012). Ageing also has a pronounced effect on sleep, including sleep quality, latency, and distribution of time within specific sleep stages (Ohayon et al., 2004). The consequences of insufficient or disrupted sleep extend beyond daily fatigue, with chronic poor sleep having far-reaching effects on cognitive performance (Alhola & Polo-Kantola, 2007) and long-term health outcomes (Medic et al., 2017).

Sleep and cognition are intricately linked, with deficits in one often predicting or contributing to deficits in the other (Nebes et al., 2009). Cognitive impairments increase with ageing and are hallmark characteristics of neurodegenerative conditions such as Alzheimer’s disease – understanding the relationship between sleep and cognitive decline is important for linking sleep and cognition with neuropathology. The interplay between sleep-cognition includes the capacity to navigate complex social and environmental challenges (Beattie et al., 2015; Simon et al., 2020), and broader impacts such as educational (Okano et al., 2019) and professional success (Barger et al., 2005; Landrigan et al., 2004; Lockley et al., 2004). The relationship between sleep and cognition is a clinical concern because both are key predictors of long-term health and the ability to function independently in society (Dew et al., 2003; McGuire et al., 2006). Poor sleep has been associated with an increased risk of developing cardiovascular disease, metabolic disorders, depression, and neurodegenerative diseases (Fang et al., 2019; Musiek & Holtzman, 2016; St-Onge et al., 2016), with accelerated cognitive decline leading to a substantially diminished quality of life, loss of independence, and a growing societal and healthcare burden, with sleep and cognition playing crucial roles in the trajectory of ageing (Scullin & Bliwise, 2015), making sleep quality a target for early interventions designed to preserve cognitive health and enhance longevity.

Sleep and cognition are served by complex neural circuits (Perrault et al., 2025). Imaging these circuits non-invasively with magnetoencephalography (MEG) has proven a powerful approach to understanding functional brain health and disease, measuring the magnetic fields generated by neural electrical activity, offering insight into the brain’s rapid, millisecond-scale oscillatory dynamics (Gross, 2019), while remaining completely non-invasive. Neural oscillations, rhythmic patterns of electrical activity in the brain, are fundamental to brain function, coordinating communication between different brain regions and supporting processes like cognition, perception, and motor control (Abubaker et al., 2021). Functional connectivity (FC), which reflects the coordinated flow of information and communication between different brain regions, is mediated by regional, time-varying changes in the amplitude or phase of oscillatory activity, and has been linked to a wide variety of cognitive processes (Ward, 2003).

Neural activity in sleep deprivation presents with excess low-frequency content in the delta and theta bands (Aeschbach et al., 1997; Cajochen et al., 1995; Finelli et al., 2000; Lian et al., 2023), which is interpreted as a sign of impaired vigilance (Del Percio et al., 2019). Notably, low-frequency activity becomes dominant in frontal brain regions; this anterior slowing is thought to reflect compensatory mechanisms to maintain cognitive performance under insufficient sleep (Goel et al., 2009; Killgore, 2010). Conversely, decreases in alpha activity occur during sleep deprivation, in anterior (Boonstra et al., 2005) and posterior regions (Del Percio et al., 2019; Lian et al., 2023; Wu et al., 2021), reflecting disruptions in relaxation and attentional processes (Abubaker et al., 2021). Prolonged wakefulness is associated with homeostatic and circadian influences, which in turn modulate neural oscillations, each with distinct spatio-temporal characteristics (Aeschbach et al., 1997). For functional connectivity, decreased coupling has been observed in the default mode network (DMN) related to sleep deprivation, with most prominent effects in the precuneus (Lian et al., 2023; Wu et al., 2021) and posterior cingulate cortex (Wang et al., 2020).

Similarly, neural oscillations are altered along the continuum of cognitive impairment, beginning with subclinical changes in cognition (López-Sanz et al., 2016), through mild cognitive impairment (MCI) (Garcés et al., 2013), and in dementia conditions such as Alzheimer’s disease (AD) (Mandal et al., 2018). In those with a genetic predisposition for AD, the prodromal stage includes neuronal hyperactivity (van Nifterick et al., 2022), with a progressive shift to oscillatory/neural ‘slowing’ which includes excess low-frequency content (Wiesman et al., 2022), with increased delta and theta activity, as well as diminished alpha and beta content (Babiloni et al., 2020; Giustiniani et al., 2023; Horvath et al., 2018). The magnitude of these effects is directly associated with cognitive deterioration (Gallego-Rudolf et al., 2024), corresponding to neurodegenerative disease stage/severity (Lejko et al., 2020). Changes in functional connectivity are also implicated in cognitive impairment, including reduced posterior alpha and beta connectivity (Babiloni et al., 2020) in dementia, as well as lower interhemispheric connectivity in delta and alpha, and reduced fronto-parietal and fronto-temporal connections in alpha (Babiloni et al., 2016). Additionally, subjective cognitive decline (SCD) has also been shown to exhibit similarly altered FC as MCI (López-Sanz et al., 2017). Given the known independent neurophysiological effects of sleep deprivation and cognitive decline, we hypothesised that poor sleep quality would be associated with neural slowing and reduced alpha and beta connectivity independent of age.

In this study, we identify a common neurophysiological profile that is associated with both sleep quality and cognitive abilities. By leveraging the large normative Cam-CAN MEG dataset, we examine key features of neural activity such as meso- and macro-scale synchronisation, including regional periodicity and network functional connectivity. We observed direct associations between sleep quality, cognitive performance, and intrinsic neurophysiological activity, which could support biomarker-driven identification of sleep and cognitive impairment, and support targeted neural interventions for restoring mental fatigue, sleep, and cognition.

## Methods

### Participants

We used data from the Cambridge Centre for Ageing and Neuroscience (Cam-CAN), a large-scale collaborative research project which collects epidemiological, cognitive, and neuroimaging data to understand ageing (https://cam-can.mrc-cbu.cam.ac.uk/dataset/) (Shafto et al., 2014). Data collection and sharing for this project was provided by CamCAN. CamCAN funding was provided by the UK Biotechnology and Biological Sciences Research Council (grant number BB/H008217/1), together with support from the UK Medical Research Council and University of Cambridge, UK. Multiple neuroimaging modalities are employed, including MEG and structural magnetic resonance imaging (MRI). T1-weighted structural images were acquired using a Magnetization Prepared RApid Gradient Echo (MPRAGE), and resting-state eyes-closed MEG data were collected using the VectorView MEG system (Elekta Neuromag, Helsinki).

Additionally, demographic, sleep quality, and cognitive measures were collected across healthy participants approximately spread across sex and age ranges. The entire dataset is composed of n=647 participants with resting-state MEG data, of which 576 passed our MEG quality assurance process (Hardy et al., 2024) and also had usable MRIs. Furthermore, 86 participants were excluded on the basis of exhibiting maximum head movement in excess of 10 millimetres during the MEG recording, resulting in a final total of n=490 participants (see **Table 1** and **Figure 1** for demographic information). In accordance with the Declaration of Helsinki all participants provided prior informed consent for the study; access to Cam-CAN was approved and granted under the terms of the Cam-CAN data use agreement. All data analyses have been approved by the relevant local ethics committee (Shafto et al., 2014).

**Figure 1:**
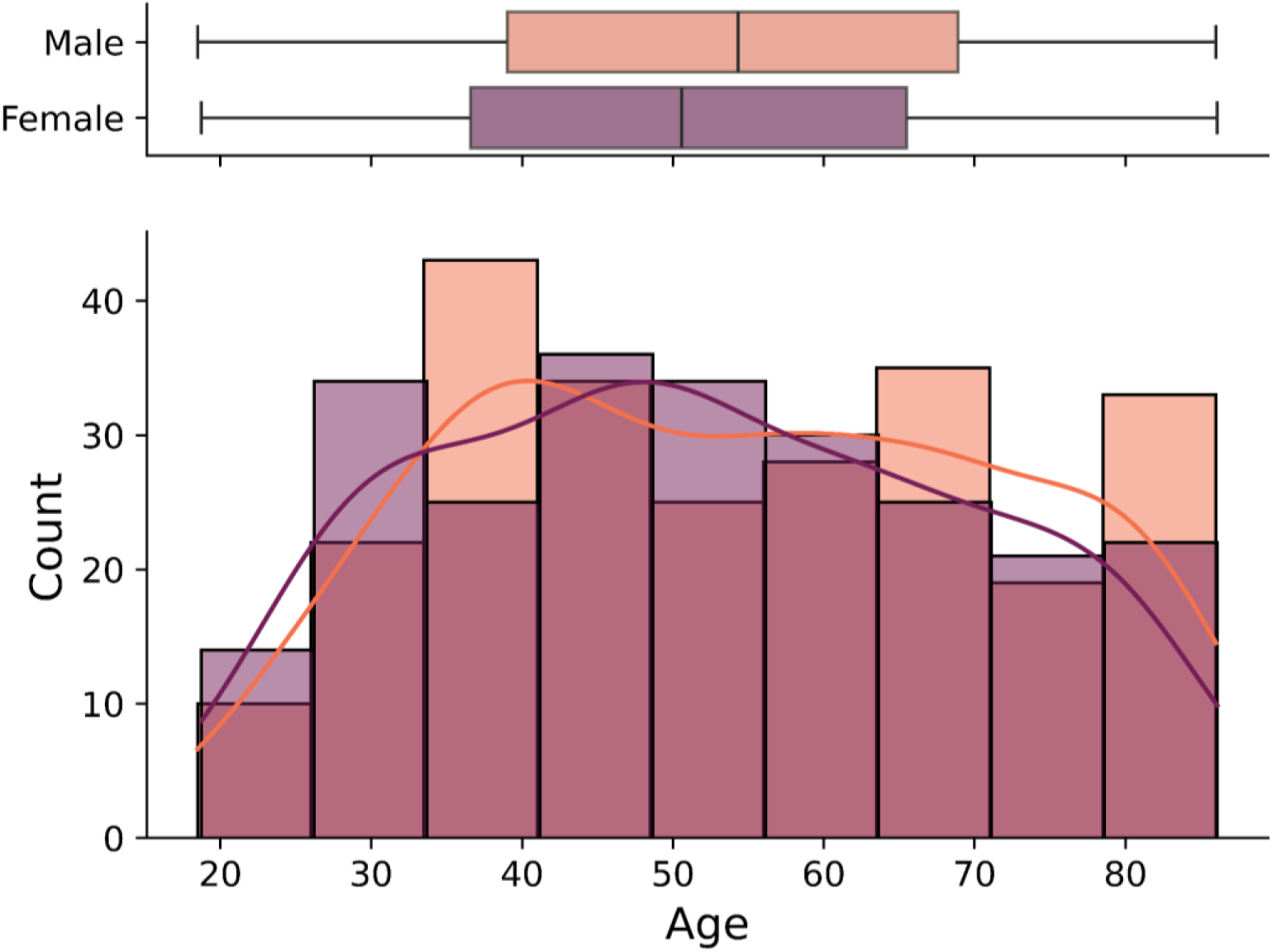
Histogram of participant age, grouped by sex.

**Table 1:**
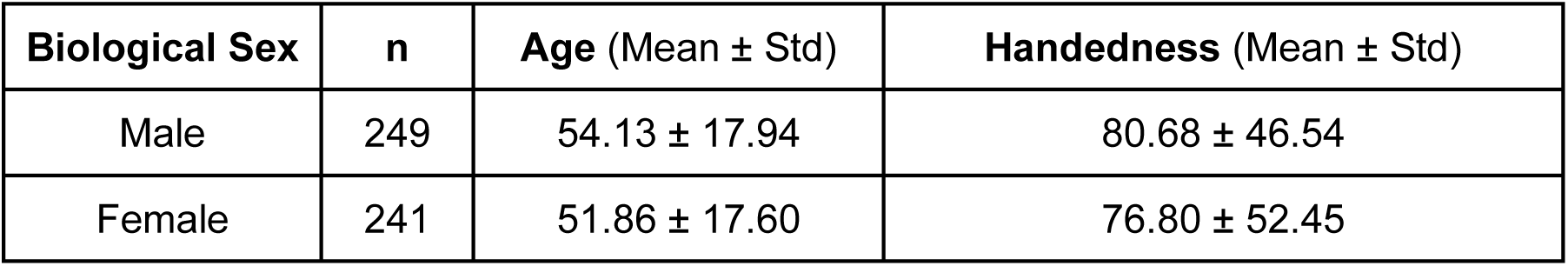
Participant demographic characteristics.

### MEG Data

Eyes-closed resting-state MEG data was collected using the VectorView MEG system (Elekta Neuromag, Helsinki) with participants in a seated position. Magnetic field data were digitised at a sampling rate of 1 kHz with a hardware bandpass filter of 0.03 - 330 Hz. To minimise environmental magnetic interference, the MaxFilter algorithm (Taulu & Simola, 2006) was applied to the filtered data with a correlation window of 10s and correlation limit of 0.98. Each participant’s scalp surface outline was measured with a Polhemus digitizer, which facilitated the co-registration of the source model to the sensor coordinate system.

Preprocessing of MEG data was performed by an automated pipeline using the MNE-Python open-source software package. Initially, the recordings were downsampled to 300 Hz and band-pass filtered between 1 and 150 Hz, with the 50 Hz power line peak removed by a notch filter with 1.5 Hz bandwidth. An additional high-pass filtered version of the data, at 110 Hz, was created to detect high-frequency muscle artefacts. Continuous head movement was tracked via fiducial coil fields.

Following filtering, the data were segmented into epochs of length 10.24s. Any epochs showing peak-to-peak amplitudes exceeding 6000 fT or containing artefactual muscle activity, determined as peak-to-peak amplitudes exceeding 2000 fT in high-frequency filtered data (110 Hz to 140 Hz) were discarded to prevent contamination by muscle artefacts or low-frequency interference. The first 25 artefact-free epochs were selected for analysis, providing 256 seconds of clean data for each participant. After filtering and epoching, the data were processed as outlined in our prior work (Hardy et al., 2024), including use of structural MRIs for co-registration.

Source-resolved data were used to derive spectral power estimates at 78 locations from the Automated Anatomical Labelling atlas for five canonical frequency bands of delta (1-3Hz), theta (4-8Hz), alpha (9-12Hz), beta (13-29Hz), and gamma (30-45Hz). Furthermore, functional connectivity was calculated using amplitude envelope connectivity at 44 seeds distributed throughout the central executive, DMN, motor, visual, and attention networks (see **Supplementary Materials**).

### Sleep Quality

The Pittsburgh Sleep Quality Index (PSQI) was used to assess participants’ sleep quality. The PSQI is a standardised, self-report questionnaire that evaluates subjective sleep quality over a one-month period (Buysse et al., 1989). It has been widely validated in both clinical and non-clinical populations, making it a highly reliable tool for identifying sleep disorders and dysfunction (Mollayeva et al., 2016). The PSQI has been linked to outcomes in physical and mental health (Hayashino et al., 2010), making it particularly relevant for research on sleep patterns and their broader implications.

The PSQI generates a global score based on seven component subscores, each reflecting a specific dimension of sleep quality; the subscores include subjective sleep quality, sleep latency, sleep duration, sleep efficiency, sleep disturbances, sleep medication, and daytime dysfunction. Each of these components is scored on a scale of 0 to 3, with higher scores indicating greater dysfunction, contributing to a global score that ranges from 0 to 21. Typically, a global score greater than 5 suggests poor sleep quality (Buysse et al., 1989). This multidimensional approach makes the PSQI particularly suitable for this study’s objectives.

### Cognitive Measures

The Addenbrooke’s Cognitive Examination - Revised (ACE-R) was used to assess participants’ cognitive functioning. The ACE-R is a widely recognised, clinically validated cognitive screening tool that is particularly sensitive to early cognitive impairments, such as those seen in dementia (Crawford et al., 2012). Its comprehensive nature and reliability make it an invaluable instrument for evaluating cognitive deficits across multiple domains (Hodges & Larner, 2017). The ACE-R’s clinical relevance is well established, as it provides a more detailed analysis of cognitive abilities compared to shorter screening tests like the Mini-Mental State Examination (MMSE) (Hodges & Larner, 2017; Pendlebury et al., 2012), with which it shares a subset of items.

The ACE-R produces a total score out of 100, which is derived from five cognitive domains, namely, attention and orientation, memory, verbal fluency, language, and visuospatial abilities. Each of these components contributes to the overall score, with higher scores indicating better cognitive performance. Scoring thresholds are typically set dynamically for each disorder to achieve the desired balance of sensitivity and specificity, for dementia, the main two thresholds are 88 (sensitivity = 0.94, specificity = 0.89) and 82 (sensitivity = 0.84, specificity = 1.0) (Mioshi et al., 2006). Additionally, the detailed domain-specific scores provide insights into which aspects of cognition may be more affected in the study population.

### Normative Modelling

To account for potential confounding variables (age, sex, and handedness) in the MEG data, normative modelling was applied using linear regression (Fraza et al., 2021; Rutherford et al., 2022). Whilst non-linear methods are often beneficial to goodness of fit, we have previously demonstrated the validity of using linear modelling of these confounding variables on the explored neurophysiological features (Hardy et al., 2024). This approach removes and accounts for the influence of sources of variance (e.g. demographic factors) on our primary MEG features: total band-limited spectral power (including periodic and aperiodic features) and functional connectivity. In each model, age, sex, and handedness were included as independent variables, while the MEG feature of interest served as the dependent variable. By fitting this model to the data, we estimated the expected value of each MEG feature based on the participants’ demographic characteristics with the residuals representing the variance unexplained by these factors, which were then used as adjusted MEG features for subsequent analysis. This regression-based normalisation allowed us to isolate the brain’s intrinsic neural features, minimising confounding effects.

### Exploratory Modelling with Partial Least Squares

The mathematics behind this approach are described fully in our previous work (Hardy et al., 2024), a brief explanation of the employed methods are as follows in this section. An exploratory partial least squares (PLS) analysis was used to model the relationship between oscillatory activity and functional connectivity (*X*) and participants’ sleep and cognitive scores (*y*). PLS is a multivariate technique that identifies latent variables, or components, which capture the shared variance between two sets of variables, in this case, MEG features and behavioural data - revealing brain-behaviour relationships.

To identify the latent structure linking *X* and *y*, a singular value decomposition (SVD) of the covariance matrix *X*^*T*^*y* was performed. This step decomposes the data into singular values and corresponding singular vectors, which represent the latent variables that capture the most variance shared between the MEG features and the sleep/cognitive scores. To determine the significance of the latent variables identified by SVD, a permutation testing procedure was used. First, we calculated the singular values from the observed data, reflecting the strength of the relationship between the MEG features and the behavioural outcomes. Then, a null distribution of singular values was generated by randomly permuting the rows (each participant) of the *y* matrix while keeping *X* fixed, effectively shuffling the correspondence between the MEG features and behavioural outcomes of each participant. This permutation procedure was repeated 10,000 times, each time recalculating the singular values. Z-Scores were generated for the singular values from the observed data by comparison against the null distribution of singular values derived from the permuted *X* and *y* matrices. P-values were calculated using the Z-Scores and the Gaussian survival function, and the results were doubled to ensure two-sided testing. Benjamini/Hochberg false discovery rate (FDR) correction (Benjamini & Hochberg, 1995) was used throughout this work, including on latent variable P-values as well as subsequent correlation tests. A latent variable was deemed significant if its observed singular value resulted in an FDR adjusted P-value below the statistical significance threshold of 0.05, indicating that the observed relationship between the MEG features and the sleep/cognitive scores was unlikely to have arisen by chance, i.e. we reject the null hypothesis that the MEG features and behavioural outcomes are unrelated (and the witnessed relationships occur by chance).

The significant latent variable(s) identified were then interpreted in terms of their contributions to both the MEG features and the sleep/cognitive scores. The weights on *X* (MEG features) and *y* (sleep and cognitive scores) provided insight into which neurophysiological and behavioural measures were most strongly associated in the latent space (X scores and y scores representing MEG and sleep/cognitive performance respectively in the latent space), thereby revealing patterns of brain-behaviour interactions.

## Results

### Poor sleep and cognitive performance are mediated by a shared neurophysiological profile

Exploratory analysis of the y matrix prior to PLS highlights the relationships between sleep scales and cognitive outcomes, as shown in **Figure 2**. PLS analysis identified one significant latent variable (p < 0.05), indicating a relationship between neural indices, and sleep and cognitive scores. The y weights for this latent variable (**Figure 3A**) illustrate that better cognitive performance—particularly in visuospatial abilities, language, fluency, and memory—was positively associated with this neurophysiological profile. In contrast, better sleep quality, as indicated by lower scores on sleep dysfunction, sleep latency, and subjective sleep quality, was negatively associated with this same latent variable. This suggests that a common neural mechanism, reflected in cortical oscillatory activity and functional connectivity, may underlie both better sleep quality and cognitive function. To this effect, **Figure 3B** shows that the neural profile is correlated with sleep-cognition outcomes (r(488)=.27, p<.001).

**Figure 2:**
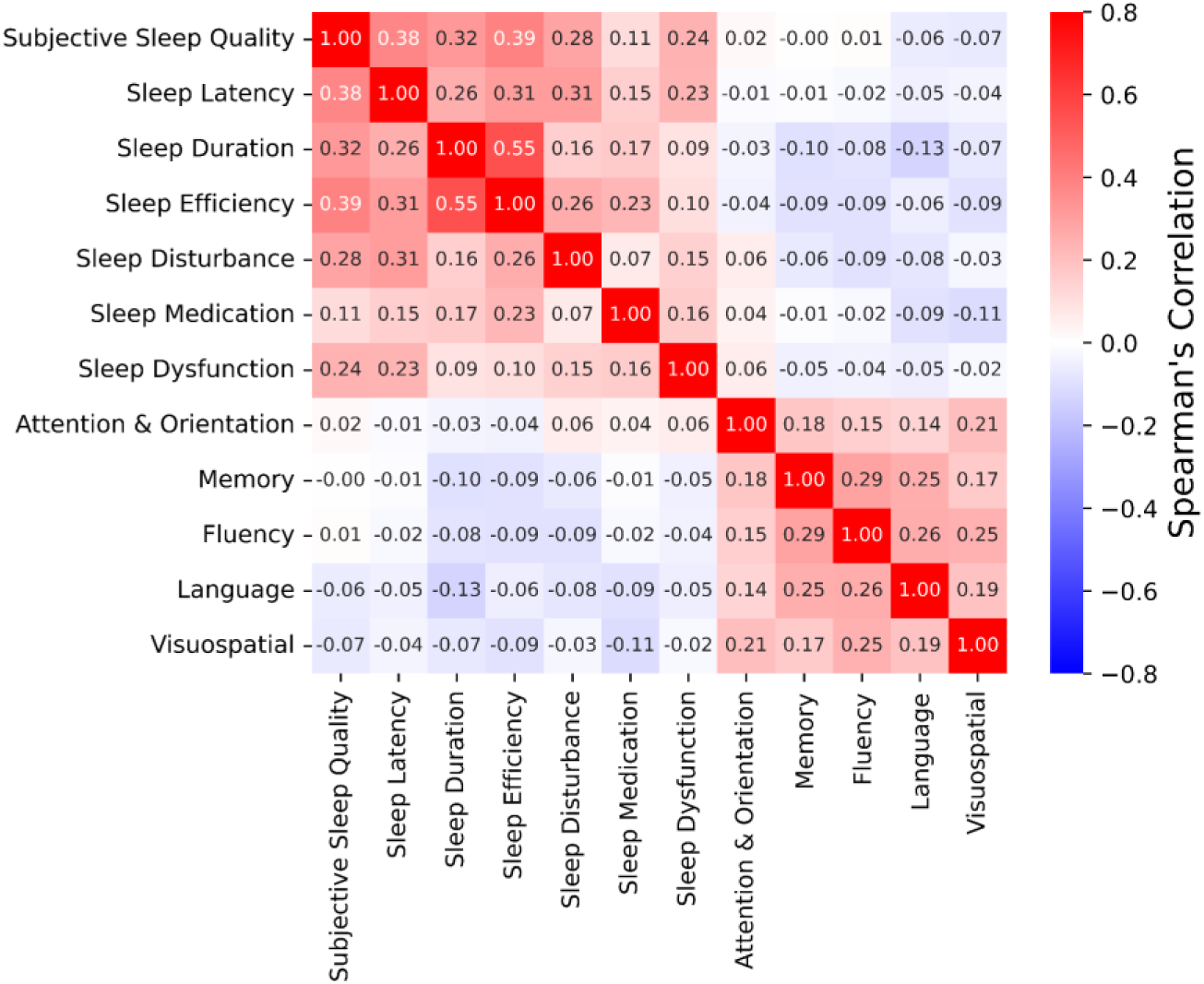
Correlation matrix highlighting the Spearman’s correlation value within and between sleep and cognitive score attributes. Dimensions of sleep generally positively correlate with one another, as do cognitive performance.

**Figure 3:**
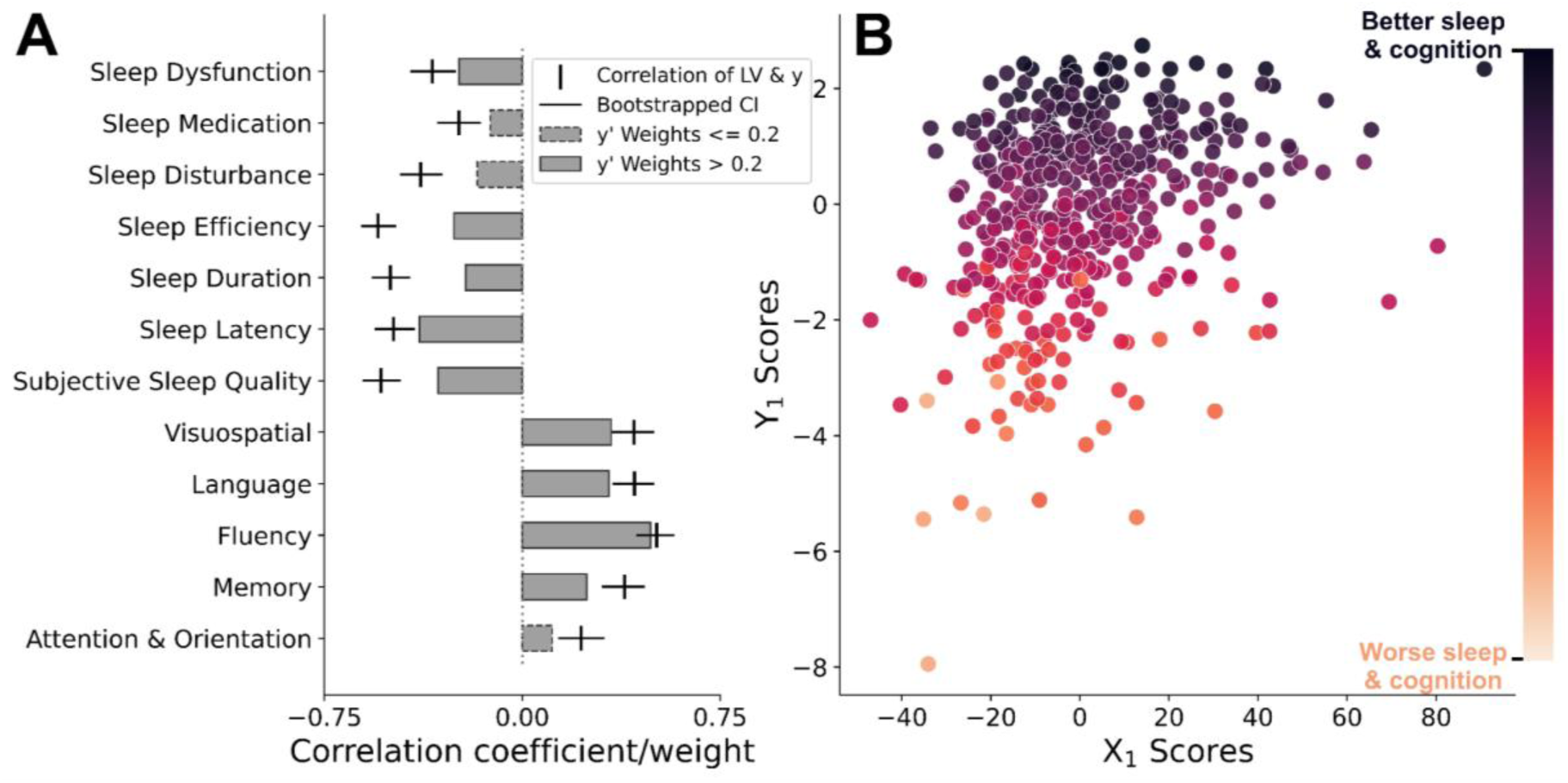
Poor sleep quality maps onto poorer cognitive performance. A) y weights and correlation coefficients between y values and the y scores. The y weights highlight the strength and direction of relations between improved sleep quality (e.g. lower scores) and better cognitive outcomes (e.g. higher scores). B) The X and y scores of the latent variable (r(488)=.27, p<.001).

The y score of the statistically significant latent variable is correlated against each of the component sub scores from the sleep and cognitive assessments. Associations between the y score and the sleep measures range between -.23 and -.54 with the strongest effects including efficiency (r(488)=-.54, p<.001), subjective quality (r(488)=-.52, p<.001), and duration (r(488)=- .50, p<.001) of sleep. Sleep medication and disturbance exhibit low correlation and y weights, highlighting that these attributes have limited effect on the latent variable. Similarly, for cognitive measures the y score produced correlations between .22 and .51, in which verbal fluency (r(488)=.51, p<.001), language (r(488)=.43, p<.001), and visuospatial (r(488)=.42, p<.001) measures are most strongly related to the latent variable.

### Frequency-specific hyper- and hypoactivity in cortical oscillations and functional connectivity are linked to deficits in sleep and cognition

Investigation of the X weights (used to project MEG data into the X scores) of the latent variable revealed a distinct spatial pattern of transpectral neural oscillatory power distributed across multiple brain regions associated with better sleep quality and cognitive outcomes (see **Figure 4**). Specifically, reductions in low-frequency power, particularly in the theta band, was observed in posterior regions, primarily parieto-occipital cortices. Similarly, lower alpha activity in fronto-parietal areas, corresponding to somatomotor cortices, and higher occipital alpha activity, was associated with better sleep and cognition.

**Figure 4:**
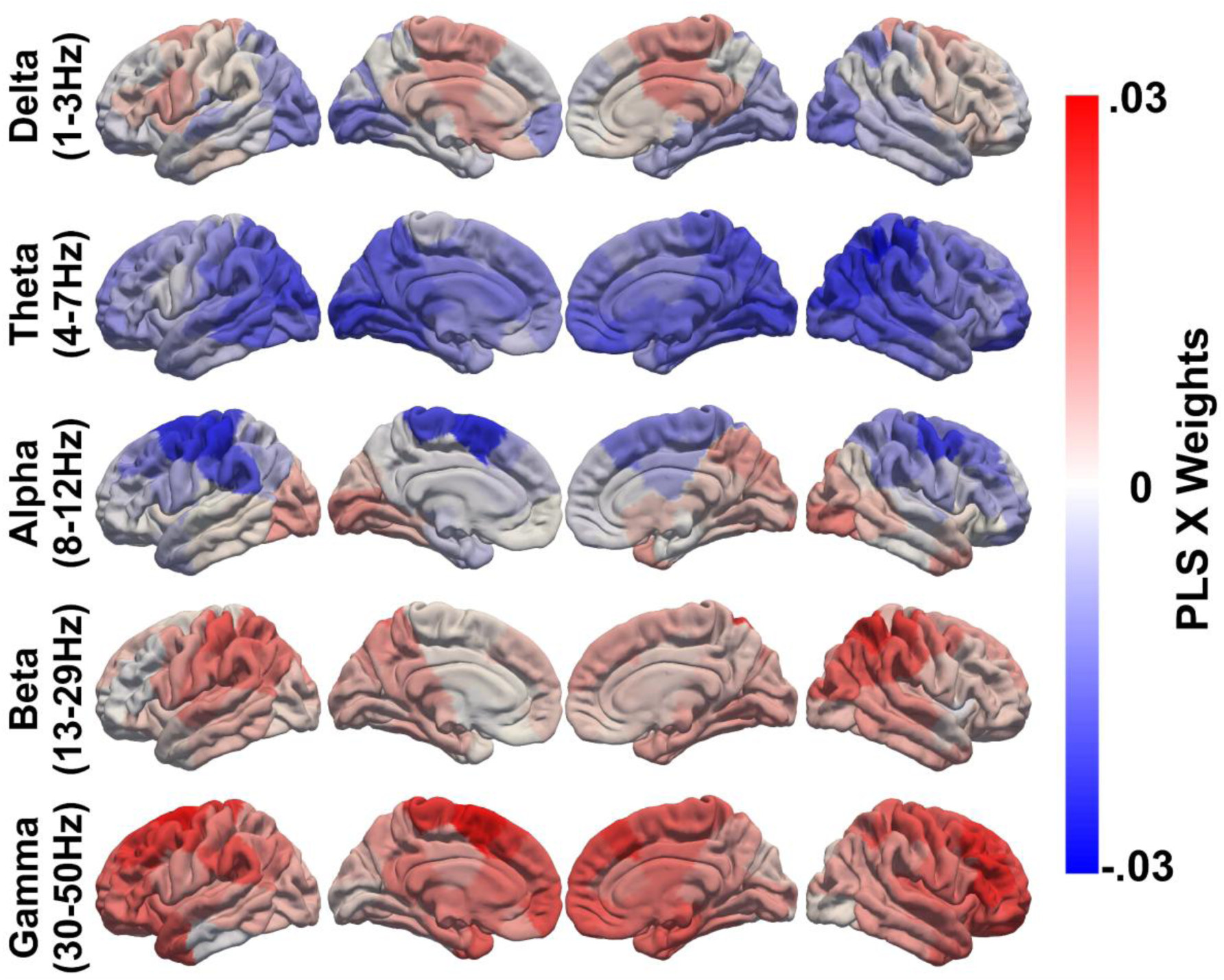
Spectral power weights for the statistically significant latent variable are shown across frequency bands of interest. The results reveal a dual pattern, with ‘better’ sleep and cognition associated with reduced slow-wave power and enhanced gamma activity. These effects are spatially specific: negative weights are observed for occipital delta, global theta, and fronto-parietal alpha bands, whereas positive weights emerge in occipital alpha, parietal beta, and fronto-parietal gamma regions.

In contrast, elevated higher frequency activity associated with this profile was observed for beta in fronto-parietal areas, and frontal and midline regions for gamma activity. These findings suggest that positive sleep and cognitive outcomes are marked by a general balance of reduced low-frequency activity in posterior brain regions and heightened high-frequency activity in parietal and fronto-parietal areas.

Association between the X scores of the significant latent variable and X values representing spectral power and functional connectivity were established via correlation. Within spectral power features the correlations ranged between -.55 and .39, with the results reflecting the same region and band-wise relationships present in the PLS X weights. Delta and theta exhibit correlations in the ranges -.19 to .05 and -.55 to -.19, respectively, including 21 and 78 of the 78 regions investigated proving statistically significant after FDR correction. The statistically significant correlations within the delta band were negatively signed (M=-.14, SD=.03) and focused in the occipital lobe, as well as posterior regions of the temporal and parietal lobes. Similarly, the strongest theta correlations are present in the posterior parietal cortex (with a maximum of r(488)=-.55, p=.004). Alpha produced the fewest band-wise significant correlations at 16 of 78, with negative associations within the dorsolateral and dorsomedial prefrontal and premotor regions, and the motor cortex. Both beta and gamma exhibit widespread positive correlations in the ranges .09 to .39 and .04 to .21, with 76 and 70 of 78 nodes being statistically significant respectively. Higher association is present bilaterally in beta across parietal and posterior temporal lobe regions, gamma, on the other hand, shows diffuse whole brain correlation. Visualisation of these findings can be found in the **Supplementary Materials, Figures SM1-SM5**.

For functional connectivity, widespread intra- and inter-network reductions in delta and theta coupling were present in the X weights, most prominently in theta (as shown in **Figure 5**), reflecting a dampening of slow-wave interactions across large-scale brain networks. Conversely, alpha and beta connectivity - the dominant mode of functional coupling in electrophysiological networks during rest - exhibited decreased connectivity within and between network connections. Gamma presents a mix of bidirectional edgewise connectivity relationships, with some regions displaying associations with sleep and cognition that involve increased coupling. In contrast, others exhibit reduced coupling, suggesting a more complex modulation of high-frequency neural communication in relation to the broader functional architecture of the brain.

**Figure 5:**
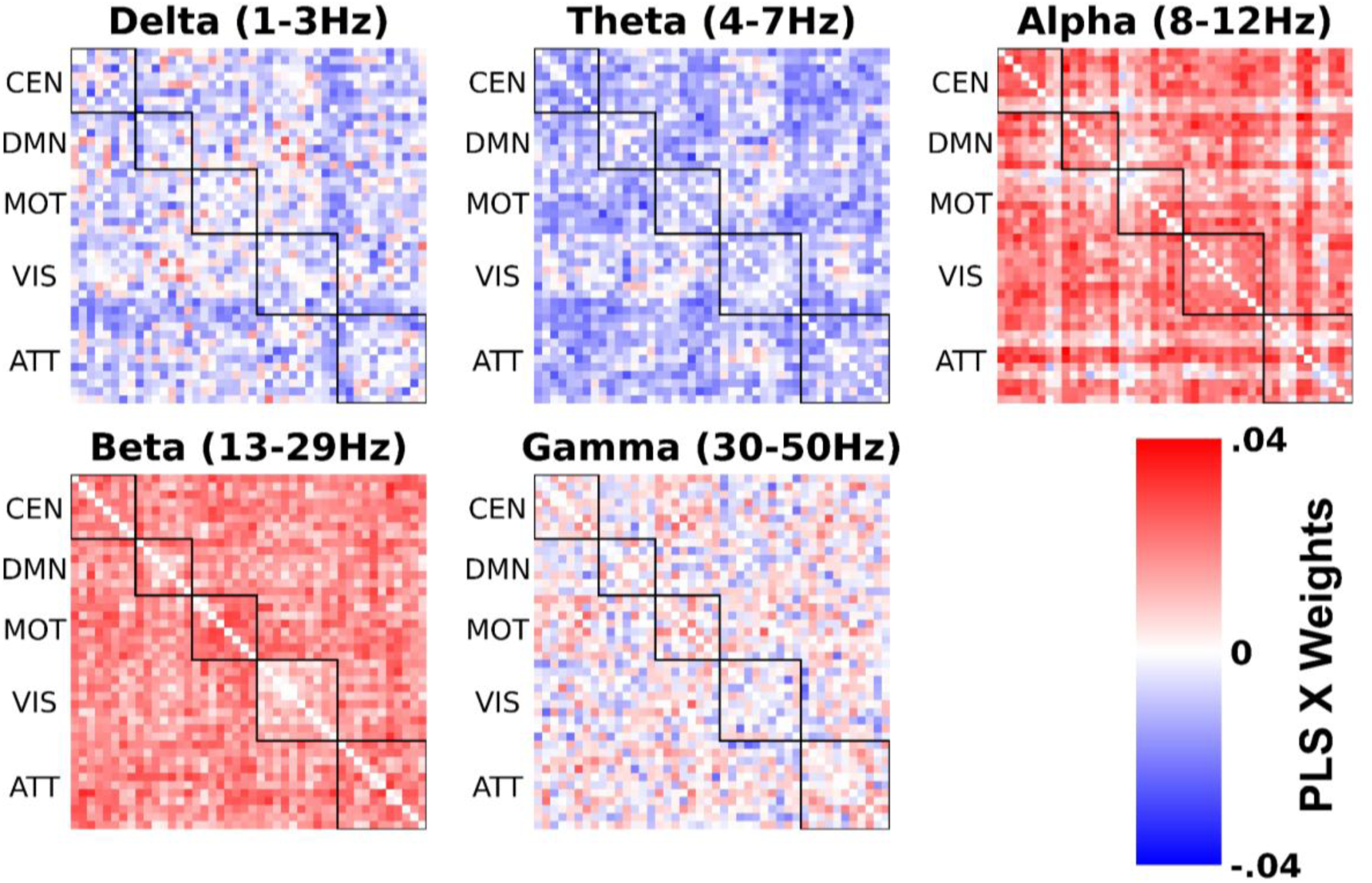
Sleep-cognition score map onto intrinsic network and between network functional connectivity across frequency ranges. The value of the X weights for Delta (1-3Hz), Theta (4-7Hz), Alpha (8-12Hz), Beta (13-29Hz), and Gamma (30-50Hz) functional connectivity for the statistically significant latent variable are shown via colour in matrix format. Each row and column corresponds to an AAL atlas location, with network nodes placed adjacently, as highlighted by the bounding boxes and network labels. Nodes from the central executive (CEN), default mode (DMN), motor (MOT), visual (VIS), and attention (ATT) networks are used. Diffuse decreased connectivity in delta and particularly theta, alongside increased alpha and beta connectivity, is associated with desirable sleep-cognition. Gamma connectivity also presents bidirectional changes in both intra and inter network connections.

Correlation between functional connectivity values and the X scores of the latent variable demonstrated clear band and network effects, including limited bidirectional association in delta connectivity (-.11 to .13), with only 24 of 946 node pairs proving significant after FDR correction. Theta connectivity produced correlations in the range -.21 to .11 to the X scores, with 269 node pairs exhibiting statistically significant association, and the strongest absolute correlations being negative within the motor network and between the central executive, motor, and attention networks. Alpha and beta functional connectivity was correlated with X scores the most prominently, with 945 and 946 statistically significant correlations respectively. These statistically significant correlations for both alpha (M=.42, SD=.12) and beta (M=.48, SD=.08) are widespread throughout the connectome, however, the top 10% of strongest correlations in both bands highlighted inter-network relationships between the central executive, default mode, visual, and attention networks. Finally, gamma connectivity exhibited primarily positive correlations in the range -.08 to .28, with 527 statistically significant connections. Analysis of the strongest correlations in gamma again highlights inter network relationships between central executive, motor, and attention networks. The correlation of functional connectivity to X scores are visualised in the **Supplementary Materials, Figures SM6-SM10**.

## Discussion

This study identifies a latent factor linking both sleep quality and cognitive performance to distinct patterns of intrinsic neurophysiological activity measured with MEG. After accounting for confounding variables, we found that increased low-frequency power, primarily in the theta band, together with decreased high-frequency power in the beta and gamma bands, was associated with poorer sleep quality and lower cognitive function across multiple domains. These spectral changes were accompanied by similar alterations in functional connectivity, predominantly within the alpha and beta bands. Overall, our findings support the view that neural slowing and interregional decoupling are key signatures of poor sleep and cognitive outcomes.

### Low-frequency activity and poor sleep-cognition outcomes

The neurophysiology of the sleep-cognition axis identified in our study exhibited a spectral profile consistent with the well-established phenomenon of neural slowing. Analysis of the spectral power X-weights revealed a clear frequency shift: poorer sleep quality and lower cognitive performance were associated with increased power in low-frequency bands, particularly delta and theta, accompanied by reduced power in higher-frequency beta and gamma bands. This characteristic pattern is a recognised electrophysiological signature previously linked to disrupted brain function and cognitive impairment.

Our findings are consistent with the literature on sleep deprivation (Hubbard et al., 2020), mental fatigue (Tran et al., 2020), and impaired cognitive abilities (Harmony, 2013). Similar to acute sleep deprivation studies, we observed increases in low-frequency power, particularly posterior theta (Wascher et al., 2014) and reductions in high-frequency power in beta and gamma associated with poor sleep and cognitive outcomes. Elevated low-frequency posterior activity–regions critical for visual processing and multisensory integration–being related to poor sleep and cognition replicates existing findings in acute sleep deprivation (Del Percio et al., 2019). Frontal regions exhibit increases in low frequency activity under sleep deprivation, and we found this to be the case in theta but not delta (**Figure 4**). Studies have reported greater slow wave activity in parietal and temporal areas in delta and theta (Cajochen et al., 1999) but observed frontal weights in delta and theta were not of greater magnitude than the surrounding regions.

Increased delta has been associated with diminished states of consciousness, including sleep, brain injury, coma, and anaesthesia (Frohlich et al., 2021). While delta is still present during conscious states, cortical deactivation during cortical ‘down states’ (i.e low amplitude phases of slow oscillations) is known to be related to reduced consciousness (Iwane et al., 2023). Delta generation is linked to thalamocortical mechanisms that periodically suppress cortical information processing and gate sensory input, leading to widespread cortical deactivation (Massimini et al., 2024). This activity during wakefulness in individuals with poor sleep perhaps signifies transient intrusions of sleep-like activity into the awake brain, a phenomenon sometimes referred to as ‘local sleep’, or reflect a more generalised state of mental fatigue and reduced cortical arousal. This is more likely to occur at rest, especially in eyes-closed resting-state. These data suggest that the brain of a poor sleeper may be operating in a state of diminished capacity for information processing.

Conversely, the ‘up states’ of slow oscillations contain increased beta and gamma activity, commonly accepted signatures of consciousness – the information-processing state. The signature of neural slowing here bears a resemblance to the electrophysiological markers consistently reported in patients along the Alzheimer’s disease (AD) continuum (Babiloni et al., 2020). The magnitude of this slowing effect has been shown to directly predict the severity of cognitive impairment across multiple domains, including attention, language, and processing speed (Wiesman et al., 2022). Furthermore, the spatial pattern of our neural slowing results closely aligns with findings from the Alzheimer’s disease (AD) literature. In particular, the increase in posterior theta activity among individuals with poorer sleep and cognition mirrors results from acute sleep deprivation studies and AD research, where posterior cortical slowing has been directly linked to regional amyloid-β burden (Wiesman et al., 2022). The presence of a similar signature in our non-clinical cohort suggests that poor sleep may contribute to alterations in brain spectral content associated with cognitive vulnerability, suggesting sleep-related mechanisms play an active role in shaping neural function (Dew et al., 2003).

### Frequency-Dependent Dysconnectivity and the Sleep-Cognition Axis

Mirroring the alterations in spectral power, poor sleep and cognition were also marked by changes in the brain’s functional network architecture. Specifically, poorer outcomes were associated with reduced alpha-beta coupling alongside increased connectivity in the delta-theta bands, reflecting a dual pattern of network dysregulation. Our connectivity results parallel observations in the dementia literature, where reduced alpha-beta connectivity is central to the “dysconnection syndrome” hypothesis of Alzheimer’s disease (Delbeuck et al., 2003). Alpha and beta oscillations enable efficient network communication, with alpha connectivity critical for long-range hub interactions and higher-order cognitive functions like attention, working memory, and executive control (Garcés et al., 2013). A systematic review of EEG studies found that reduced alpha-band connectivity is one of the most consistent findings in both MCI and AD compared to healthy controls (Babiloni et al., 2025; Bhattacharya et al., 2011). Our alpha-beta findings link with this, suggesting that sleep disturbances may induce a pattern of functional dysconnection resembling that seen in neurodegeneration.

We also report concurrent increases in delta and theta band connectivity suggesting that sleep outcomes influence network reorganisation in a fundamental manner: while high-frequency networks are becoming less integrated, low-frequency networks are becoming hypersynchronised. Low-frequency oscillations are linked to low arousal, cortical inhibition, and sleep. Increased connectivity in these bands during wakefulness likely reflects widespread, non-specific synchronisation that disrupts targeted, information-rich communication needed for complex cognition. In individuals with poor sleep and cognition, the brain appears to trade efficient high-frequency signaling for global low-frequency synchrony, representing a shift in network state rather than simple dysconnection. Bidirectional changes in gamma connectivity may reflect a mix of compensatory activity and pathological processes. Future work could utilise graph theoretical and network analyses to quantify the efficiency of these network changes following poor sleep. More generally, the dual signature of frequency-dependent connectivity changes reported here could serve as a multipurpose biomarker of neural functioning. It captures both the loss of healthy, efficient network function and the emergence of pathological, maladaptive network activity. This pattern of dysconnectivity and altered low-frequency coupling has also been implicated in other neuropsychiatric disorders characterised by cognitive deficits, such as schizophrenia (Alamian et al., 2020), suggesting it may represent a transdiagnostic marker of cognitive impairment. Future diagnostic tools could be developed to detect this specific pattern of network reorganisation, potentially identifying individuals at risk for cognitive decline long before the manifestation of overt clinical symptoms and providing a quantifiable target for therapeutic interventions.

### Brain-Behaviour Relationships

Another consideration arising from our results is the interplay between the stable dynamics of neural activity and the sleep and cognitive traits measured. The PSQI assesses sleep quality over the preceding month, capturing a relatively recent behavioural state (Buysse et al., 1989). Cognitive scores, likewise, represent a snapshot of current functional ability. In contrast, resting-state neural activity, particularly features like intrinsic functional connectivity, is often considered a stable, trait-like characteristic of an individual’s brain, which has been termed a ‘neurophysiological fingerprint’ (da Silva Castanheira et al., 2021; Shehzad et al., 2009). The strong association found between these measures implies that a behavioral pattern spanning just one month is coupled with the brain’s intrinsic, and therefore more stable, operating characteristics. This linkage suggests that even transient periods of poor sleep are either sufficient to induce a measurable shift in the brain’s fundamental activity or, conversely, that a pre-existing, less optimal brain state predisposes an individual to experience poorer sleep and cognitive function. Future work is required to establish how many nights of poor sleep are required to shift the MEG frequency spectrum towards a neural slowing state.

### Limitations

This study includes a number of limitations. First, it uses a cross-sectional design; with longitudinal data we could track changes in brain function linked to changes in sleep and cognition. Although our cross-sectional data precludes causal inference, our study shows that subjective sleep quality is linked to measurable and functionally relevant alterations in the brain’s intrinsic activity. Secondly, the sleep quality measures are all subjective self-report, rather than objective physiological data, such as metrics including actigraphy or polysomnography, etc. Finally, the potential cultural or demographic biases inherent in the Cam-CAN data.

### Conclusion

This study shows how sleep and cognition are intimately linked to the brain’s intrinsic neurophysiological activity. Using partial least squares analysis on a large, age-diverse MEG dataset with comprehensive cognitive assessments, we identified a latent factor connecting poorer sleep and lower cognitive performance to increased low-frequency (theta) and decreased high-frequency (beta-gamma) power, alongside altered alpha-beta connectivity. By examining spatially resolved, source-level oscillations, this study suggests that neural slowing and interregional decoupling underpin the sleep-cognition relationship. These findings advance our understanding of sleep-related cognitive variation and highlight the potential of MEG-derived oscillatory markers for early detection and the development of targeted, sleep-based interventions to mitigate cognitive decline.

## Data & Code Availability

Raw data is available from:

- Cambridge Centre for Ageing and Neuroscience (Cam-CAN): https://camcan-archive.mrc-cbu.cam.ac.uk/dataaccess/

Code was used from:

- Python: https://www.python.org/

Analysis:

- MNE: https://mne.tools/stable/index.html
- numpy: https://numpy.org/
- pandas: https://pandas.pydata.org/
- scipy: https://scipy.org/
- sklearn: https://scikit-learn.org/stable/index.html
- statsmodels: https://www.statsmodels.org/stable/index.html

Visualisation:

matplotlib: https://matplotlib.org/
pyvista: https://docs.pyvista.org/version/stable/
seaborn: https://seaborn.pydata.org/

## Author contributions

SH: Conceptualisation, Data curation, Formal analysis, Investigation, Methodology, Visualisation, Writing – original draft, Writing – review & editing.

GR: Conceptualisation, Data curation, Formal analysis, Investigation, Methodology, Visualisation, Writing – original draft, Writing – review & editing.

GO: Writing – original draft, Writing – review & editing.

VB: Writing – original draft, Writing – review & editing.

YP: Writing – original draft, Writing – review & editing.

ACR: Writing – original draft, Writing – review & editing.

RAS: Writing – original draft, Writing – review & editing.

BTD: Conceptualisation, Investigation, Methodology, Project administration, Supervision, Writing – original draft, Writing – review & editing

