## Supplementary Materials for "Intrinsic electrophysiological activity maps a latent dimension of poor sleep quality and reduced cognitive performance: a magnetoencephalography study using Cam-CAN data"

#### Network Nodes Definitions

##### Central Executive Network

RIPS [25, -62, 53]
RVV [36, -62, 0]
LVV [-44, -60, -6]
RSMG [32, -38, 38]
RSLOC [26, -64, 54]
LSLOC [-26, -60, 52]
RFEF [28, -4, 58]
LFEF [-26, -8, 54]

##### Default Mode Network

LAG [-43, -76, 35]
RAG [51, -64, 32]
PCC [-3, -54, 31]
vMPFC [-2, 51, 2]
dMPFC [-13, 52, 23]
RMPFC [2, 53, 24]
LITG [-57, -25, -17]

##### Motor network

Precentral_L [-39.0, -7.0, 50.0]
Precentral_R [41.0, -10.0, 51.0]
Postcentral_L [-43.0, -24.0, 47.0]
Postcentral_R [41.0, -27.0, 51.0]
Parietal_Sup_L [-24.0, -61.0, 58.0]
Parietal_Sup_R [26.0, -60.0, 61.0]
Parietal_Inf_L [-43.0, -47.0, 45.0]
Parietal_Inf_R [46.0, -48.0, 48.0]

##### Attention network

RSMG [52, -48, 28]
RFEF [30, -13, 53]
LFEF [-26, -12, 53]
LpIPS [-25, -67, 48]
RpIPS [23, -69, 49]

LMT [-43, -72, -8]
RMT [42, -70, -11]
RMFG [41, 17, 31]
RPCS [41, 2, 50]
RSTG [58, -48, 10]
RVFC [40, 21, -4]

#### Visual network

LV1 [-3, -101, -1]
RV1 [11, -88, -4]
LV2d [-8, -99, 7]
RV2d [14, -96, 13]
LV3 [-9, -96, 13]
RV3 [20, -95, 18]
LV4 [-31, -77, -17]
RV4 [27, -71, -14]
LV7 [-23, -78, 26]
RV7 [32, -78, 25]

#### Abbreviations:

"RIPS": "Right Intra Parietal Sulcus",
"RVV": "Right Ventral Visual",
"LVV": "Left Ventral Visual",
"RSMG": "Right Supramarginal Gyrus",
"RSLOC": "Right Superior Lateral Occipital Cortex",
"LSLOC": "Left Superior Lateral Occipital Cortex",
"RFEF": "Right Frontal Eye Field",
"LFEF": "Left Frontal Eye Field",
"LAG": "Left Angular Gyrus",
"RAG": "Right Angular Gyrus",
"PCC": "Posterior Cingulate Cortex",
"vMPFC": "Ventromedial Prefrontal Cortex",
"dMPFC": "Dorsomedial Prefrontal Cortex",
"RMPFC": "Rostral Medial Prefrontal Cortex",
"LITG": "Left Inferior Temporal Gyrus",
"Precentral_L": "Precentral Left",
"Precentral_R": "Precentral Right",
"Postcentral_L": "Postcentral Left",
"Postcentral_R": "Postcentral Right",
"Parietal_Sup_L": "Parietal Superior Left",
"Parietal_Sup_R": "Parietal Superior Right",
"Parietal_Inf_L": "Parietal Inferior Left",
"Parietal_Inf_R": "Parietal Inferior Right",

"LV1": "Left Visual 1",
"RV1": "Right Visual 1",
"LV2d": "Left Visual 2 (Dorsal)",
"RV2d": "Right Visual 2 (Dorsal)",
"LV3": "Left Visual 3",
"RV3": "Right Visual 3",
"LV4": "Left Visual 4",
"RV4": "Right Visual 4",
"LV7": "Left Visual 7",
"RV7": "Right Visual 7",
"LpIPS": "Left posterior Intra-Parietal Sulcus",
"RpIPS": "Right posterior Intra-Parietal Sulcus",
"LMT": "Left Middle Temporal",
"RMT": "Right Middle Temporal",
"RMFG": "Right Middle Frontal Gyrus",
"RPCS": "Right Precentral Sulcus",
"RSTG": "Right Superior Temporal Gyrus",
"RVFC": "Right Ventro-Frontal Cortex"

### Correlations Between MEG data and X scores

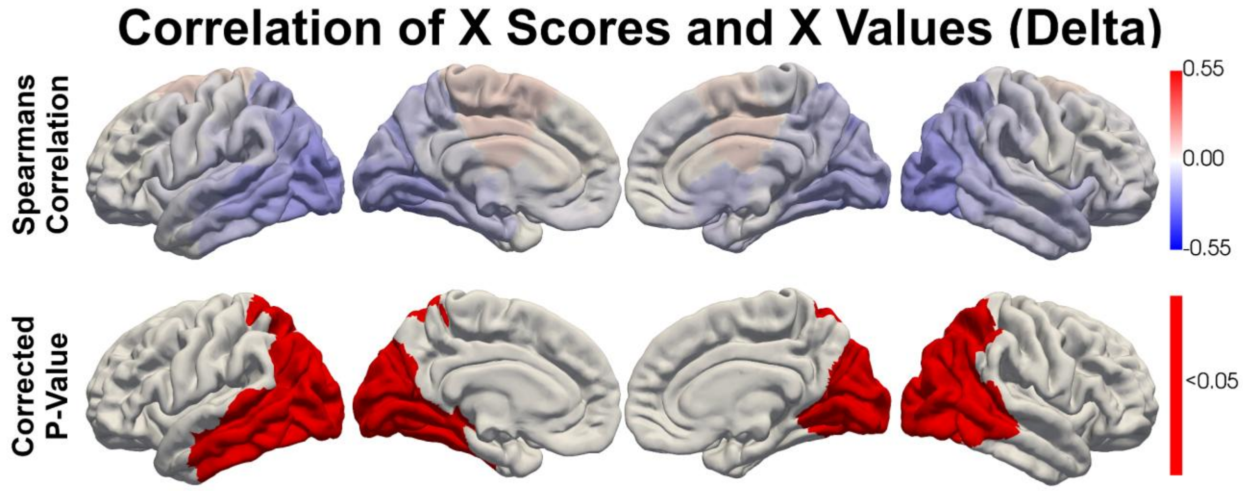


***Figure SM1:*** *Correlation between X scores and Delta (1-3Hz) spectral power, calculated via permutation testing with 1000 resamples.*

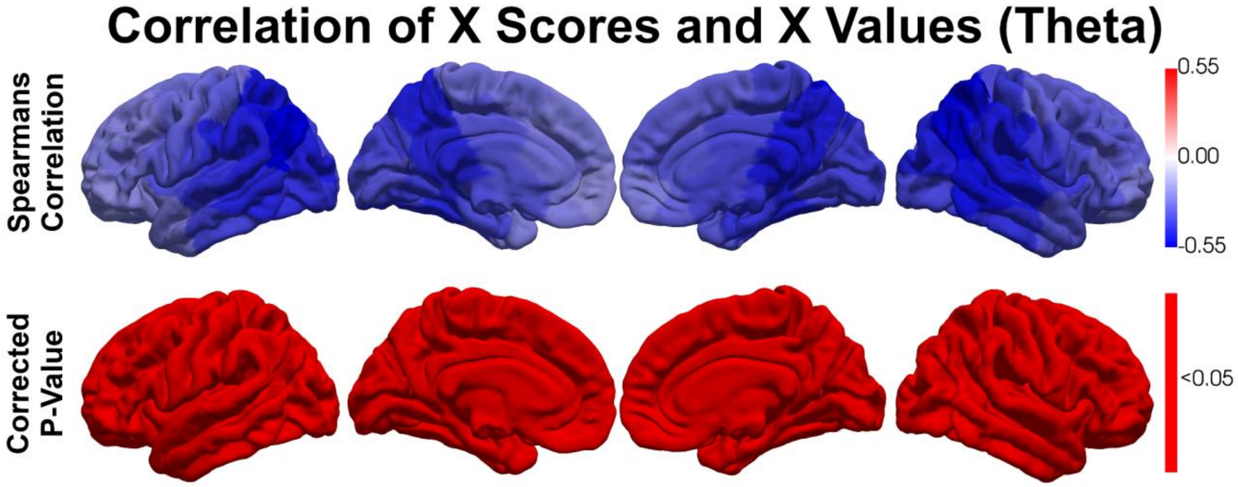


***Figure SM2:*** *Correlation between X scores and Theta (4-7Hz) spectral power, calculated via permutation testing with 1000 resamples.*

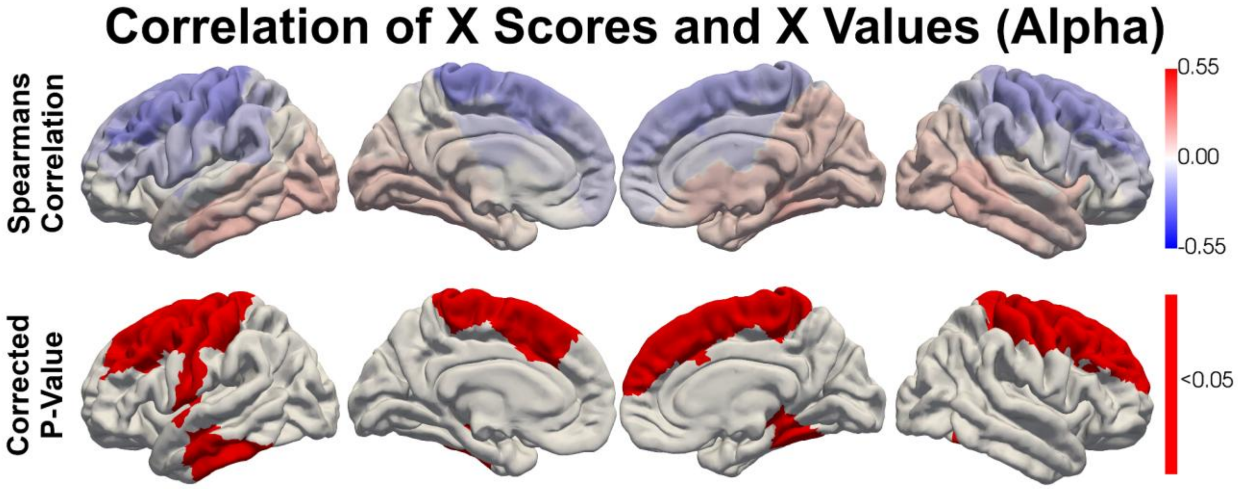


***Figure SM3:*** *Correlation between X scores and Alpha (8-12Hz) spectral power, calculated via permutation testing with 1000 resamples.*

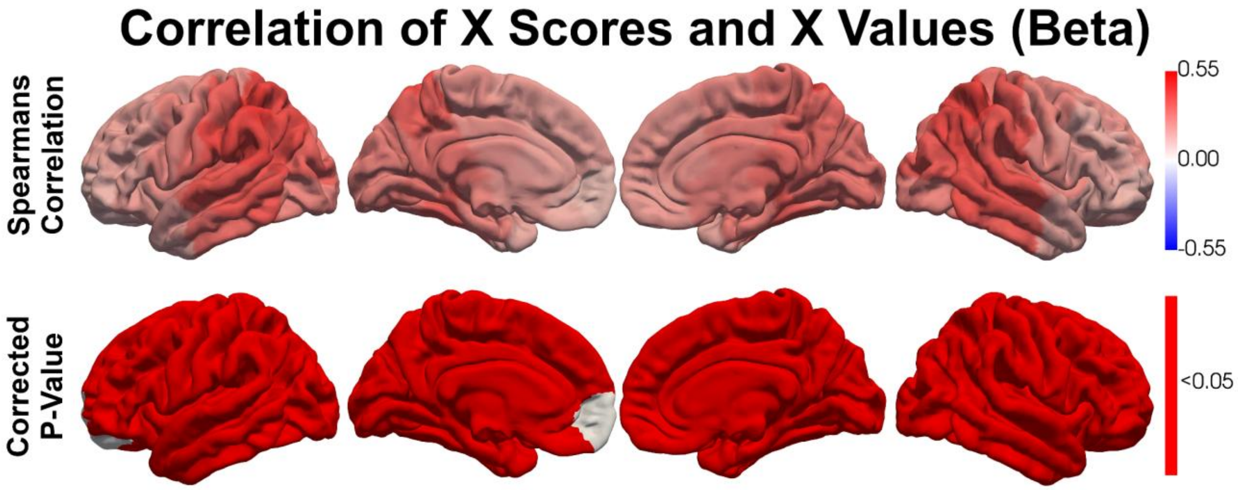


***Figure SM4:*** *Correlation between X scores and Beta (13-29Hz) spectral power, calculated via permutation testing with 1000 resamples.*

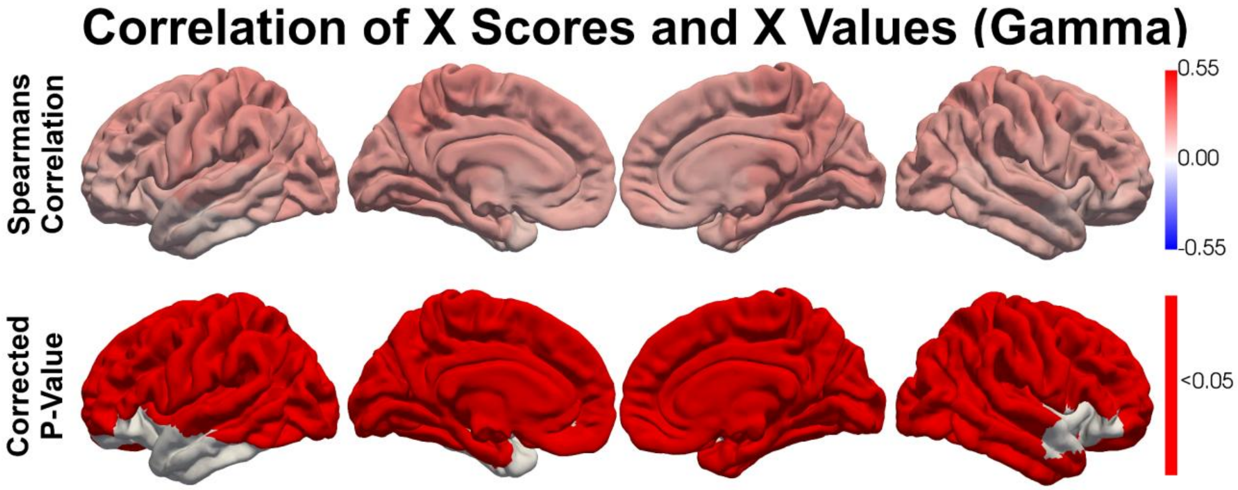


***Figure SM5:*** *Correlation between X scores and Gamma (30-50Hz) spectral power, calculated via permutation testing with 1000 resamples.*

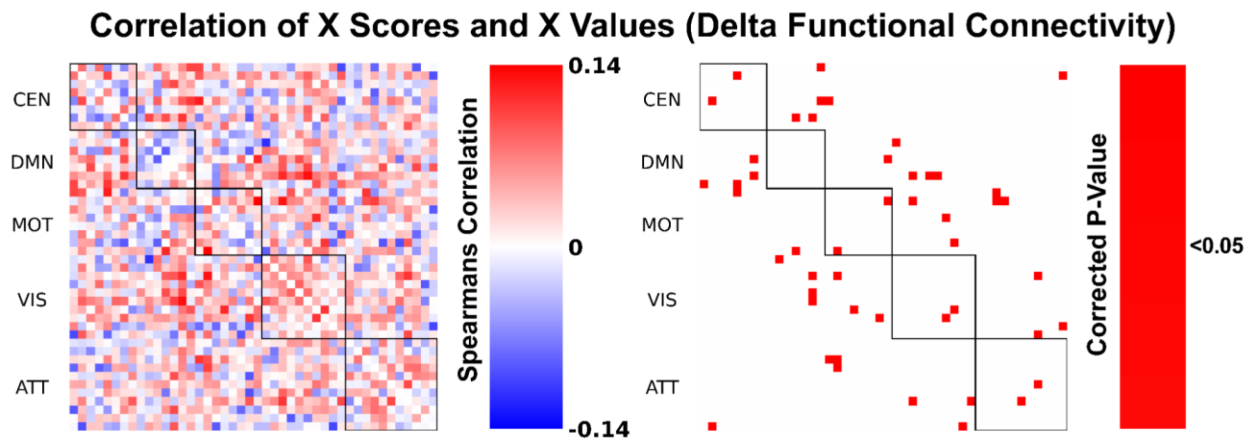


***Figure SM6:*** *Correlation between X scores and Delta (1-3Hz) functional connectivity, calculated via permutation testing with 1000 resamples.*

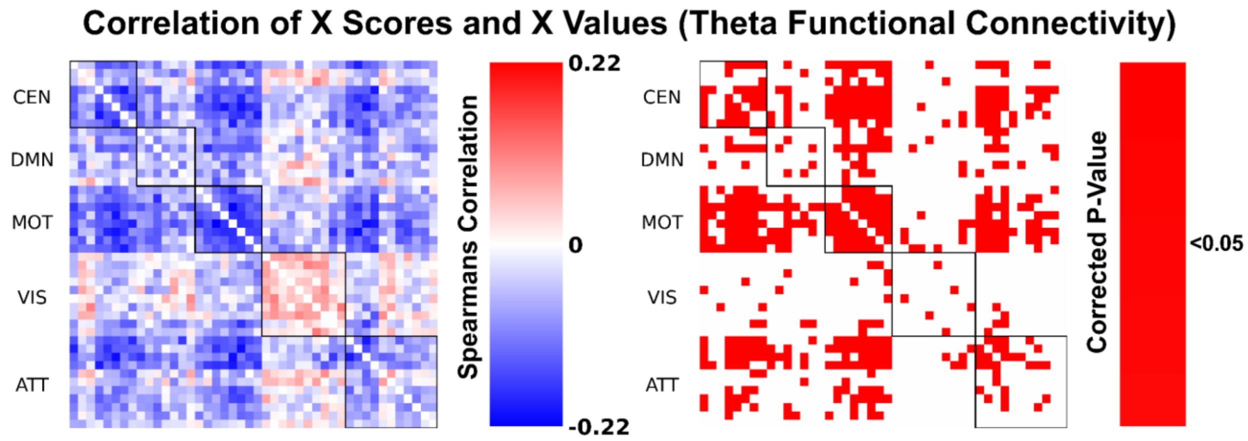


***Figure SM7:*** *Correlation between X scores and Theta (4-7Hz) functional connectivity, calculated via permutation testing with 1000 resamples.*

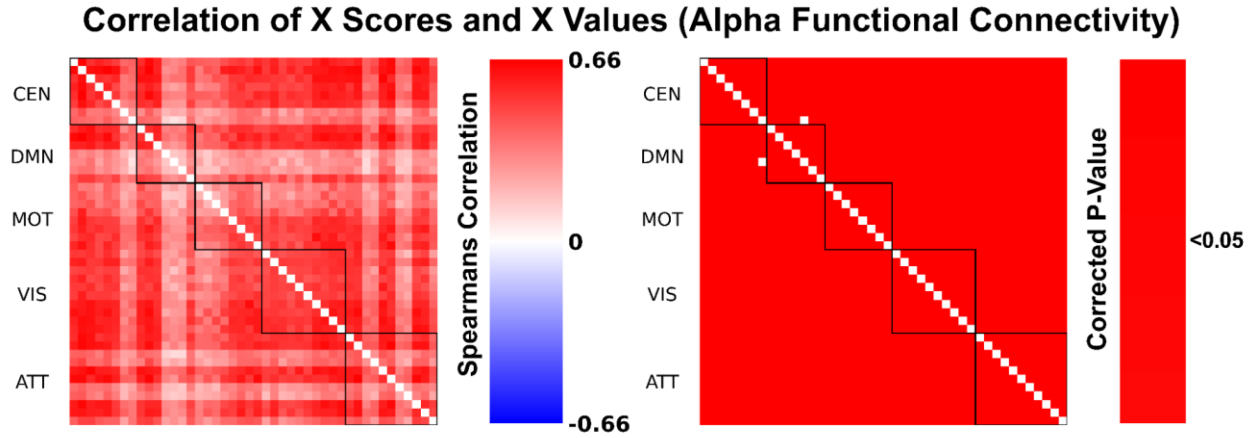


***Figure SM8:*** *Correlation between X scores and Alpha (8-12Hz) functional connectivity, calculated via permutation testing with 1000 resamples.*

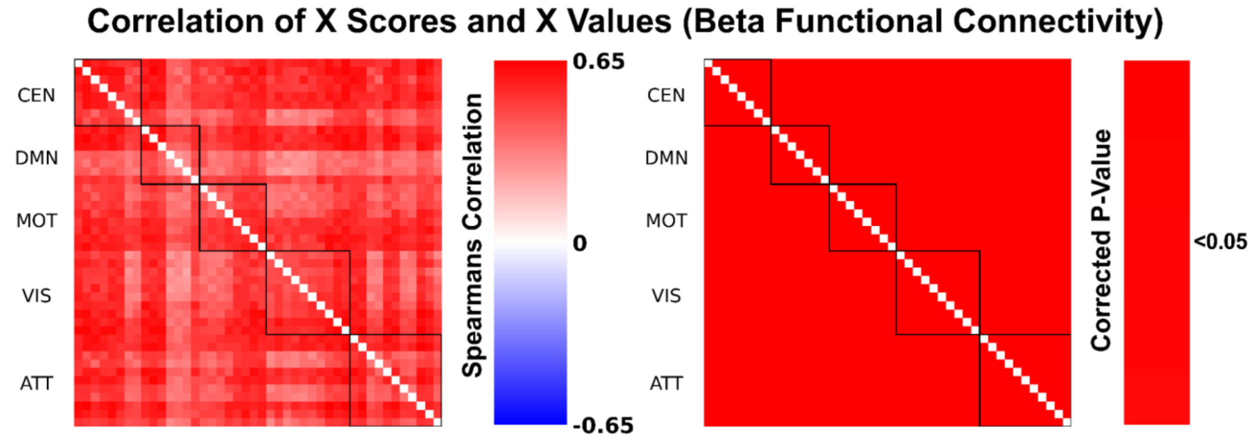


***Figure SM9:*** *Correlation between X scores and Beta (23-29Hz) functional connectivity, calculated via permutation testing with 1000 resamples.*

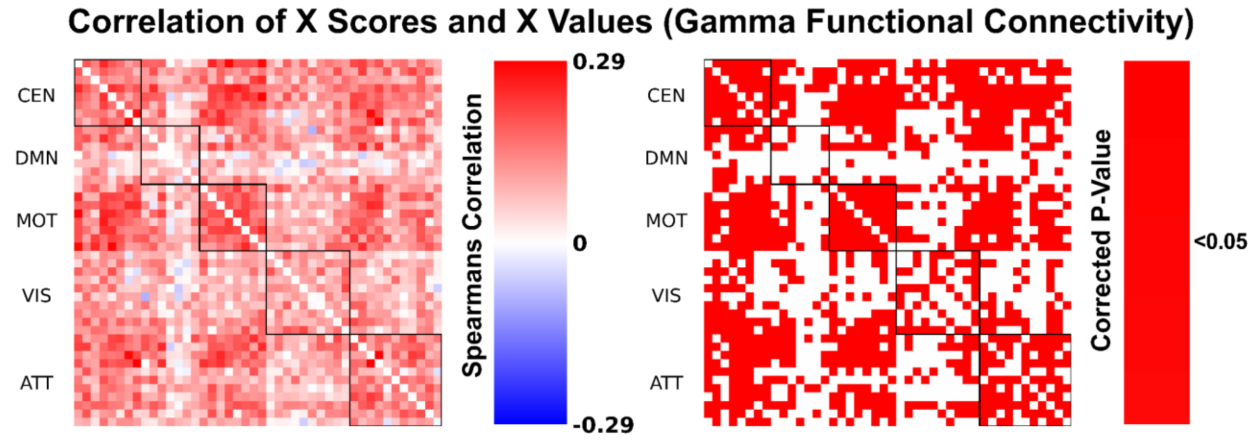


***Figure SM10:*** *Correlation between X scores and Gamma (30-50Hz) functional connectivity, calculated via permutation testing with 1000 resamples.*
